# CREB1 contributes colorectal cancer cell plasticity by regulating lncRNA CCAT1 and NF-κB pathways

**DOI:** 10.1101/2020.04.24.059162

**Authors:** Bin Li, Li-Si Zheng, Chen-Min Zhang, Qiao-Juan Huang, Yan-Hua Guo, Lu-Qin Wang, Peng Yu, Shu-Rong Liu, Qiao Lin, Yu-Xia Luo, Hui Zhou, Jian-Hua Yang, Liang-Hu Qu

**Author notes:** **Correspondence to:** Liang-Hu Qu, **e-mail**, Jian-Hua Yang, **e-mail**.

## Abstract

The CREB1 gene encodes a pleiotropic transcription factor that frequently dysregulated in cancers. CREB1 can regulates tumour cell status of proliferation or migration, however, the molecular basis for this switch involvement in cell plasticity has not been fully understood. Here, we show that knocking out CREB1 triggered a remarkable effect of epithelial-mesenchymal transition (EMT) and led to the occurrence of inhibited proliferation and enhanced motility in cancer cells. Mechanistically, CREB1-knockout cells showed arrest in the G0/G1 phase as a result of impaired CREB1-dependent transcription of CCAT1 and E2F1. Interestingly, the competition between the coactivator CBP/p300 for CREB1 and p65 leads to the activation of the NF-κB pathway in cells with CREB1 disrupted, which induces an EMT phenotype and enhances motility. These studies identified previously unknown mechanisms of CREB1 in cell plasticity via its lncRNA and protein effector pathways, revealing an important feature that should be considered in CREB1-targeted tumour therapies.

## Introduction

Cancer cells are associated with a high degree of cellular plasticity that includes molecular and phenotypic changes during tumour progression and treatment (Yuan et al., 2019). Increasing evidence suggests that cancer cells undergo phenotypic switching between growth and metastasis phenotypes in response to genomic or epigenetic alteration as well as a volatile microenvironment. Indeed, studies indicate that oncogenic genes with paradoxical functions can contribute the capacity for phenotypic plasticity. For example, *c-myc*, a well-known oncogene, can stimulate the proliferation of breast cancer cells both in culture and *in vivo*, but it can inhibit motility and invasiveness in culture and lung or liver metastases in xenografted tumours (Hong et al., 2012). Thus, the opposite role of oncogenes that can promote proliferation pathways but simultaneously suppress migration pathways might be an intrinsic mechanism of cancer cell plasticity in tumours. However, how genes with paradoxical functional role involve in cellular plasticity need to be addressed.

The oncogene *CREB1* encodes an exceptionally pleiotropic transcription factor (CREB1) that bind to promoter *cis*-regulatory elements (CREs) which are composed of TGANNTCA sequences, is ubiquitously expressed in many tissues (Impey et al., 2004, Zhang et al., 2005). Furthermore, CREB1 has been reported to regulate both coding and noncoding genes that are generally involved in the regulation of cell survival, proliferation and glucose metabolism (Lonze and Ginty, 2002, Carlezon et al., 2005, Wang et al., 2010, James et al., 2009, Fusco et al., 2016).

Elevated expression of CREB1 in multiple tumours including acute myeloid leukaemia, prostate cancer, gastric cancer and gliomas, is associated with enhanced cell proliferation (Shankar et al., 2005, Zhang et al., 2011, Rao et al., 2017, Tan et al., 2012). Paradoxically, such overexpression is dissociated from invasive and metastatic propensity, and the inhibition of CREB1 accounts for the increase of cell invasiveness (Guo et al., 2016a, Bian et al., 2016). These studies demonstrated contradictory effects of CREB1 on tumour progression, which raise the possibility that CREB1 could contribute tumour cell plasticity by switching from a proliferative to an invasive state. However, the molecular mechanisms which mediated the switch between proliferation and invasion of cancer cells are poorly understood.

Here, we investigated the impact of CREB1 on cell phenotypic plasticity and demonstrated paradoxical effects of CREB1 on cell proliferation and motility, using CRISPR/Cas9-mediated CREB1-knockout HCT116 cells. In particular, we elucidated the molecular mechanisms of CREB1 on cell plasticity by regulating lncRNA CCAT1 and NF-κB pathways. Our findings have uncovered a previously unappreciated function of CREB1 for the control of tumour cells between proliferation and metastatic states with its coding and noncoding effector pathways, which reveals a mechanism that may be important in the development of therapeutics that target CREB1.

## Results

### CREB1 deletion impairs cell proliferation and increases migration and invasion

In an effort to examine the effects of CREB1 ablation in HCT116 cells, we used CRISPR/Cas9n technology with a pair of offset sgRNAs (sgRNA-C1 and C2) complementary to opposite strands of exon 5 to knock out CREB1 (Supplementary figure 1A) (Ran et al., 2013). sgRNA-C1 and C2 were cloned into PX462, a bicistronic vector that contains the Cas9n enzyme and a programmable chimeric sgRNA. To increase the efficiency of indel mutations, the sgRNA-C2 transcript unit (U6-promoter-sgRNA-C2) was subcloned into the PX462-sgRNA-C1, resulting in the tricistronic vector PX462-C1C2 (Supplementary figure 1B). The effectiveness of PX462-C1C2 was evaluated by T7E1 nuclease assay, and the efficiency of introducing indel mutations at the exon 5 target site reached up to 90% (Supplementary figure 1C). Then, we performed a serial dilution to isolate single CREB1 knockout clones. Twenty-two clones were probed for the T7E1 nuclease assay, and eight clones were selected for additional Western blot and Sanger sequencing analyses. Compared to the unedited HCT116 cells, there was a clone (referred to as KO-CREB1) that showed a complete absence of CREB1 expression (Supplementary figure 1D and 1E).

To further evaluate whether the isolated clone was functionally CREB1 deficient, CREB1 signaling pathway activity and CREB1-associated genes (including CREB family genes and coactivators) were measured. As expected, the CREB1 signaling pathway was repressed in response to CREB1 depletion, as indicated by the significant decreased in CREB1 reporter activities and in the expression of the CREB1 target genes NR4A1, CYP19A1, BRCA1, BCL2 and cFos (Figure 1A and 1B) (Lemberger et al., 2008, Michael et al., 1997, Ghosh et al., 2008, Wilson et al., 1996, Berkowitz et al., 1989, Lee et al., 1995). Disruption of CREB1 did not affect the transcription of ATF1, CRTC1, CBP and EP300, whereas there was an approximately 4-fold increase in CREM and a marginal decrease in CREB1 (Figure 1C) (Hummler et al., 1994). Collectively, these data confirm the generation of KO-CREB1 cells using the CRISPR/Cas9n system.

**Figure 1.**
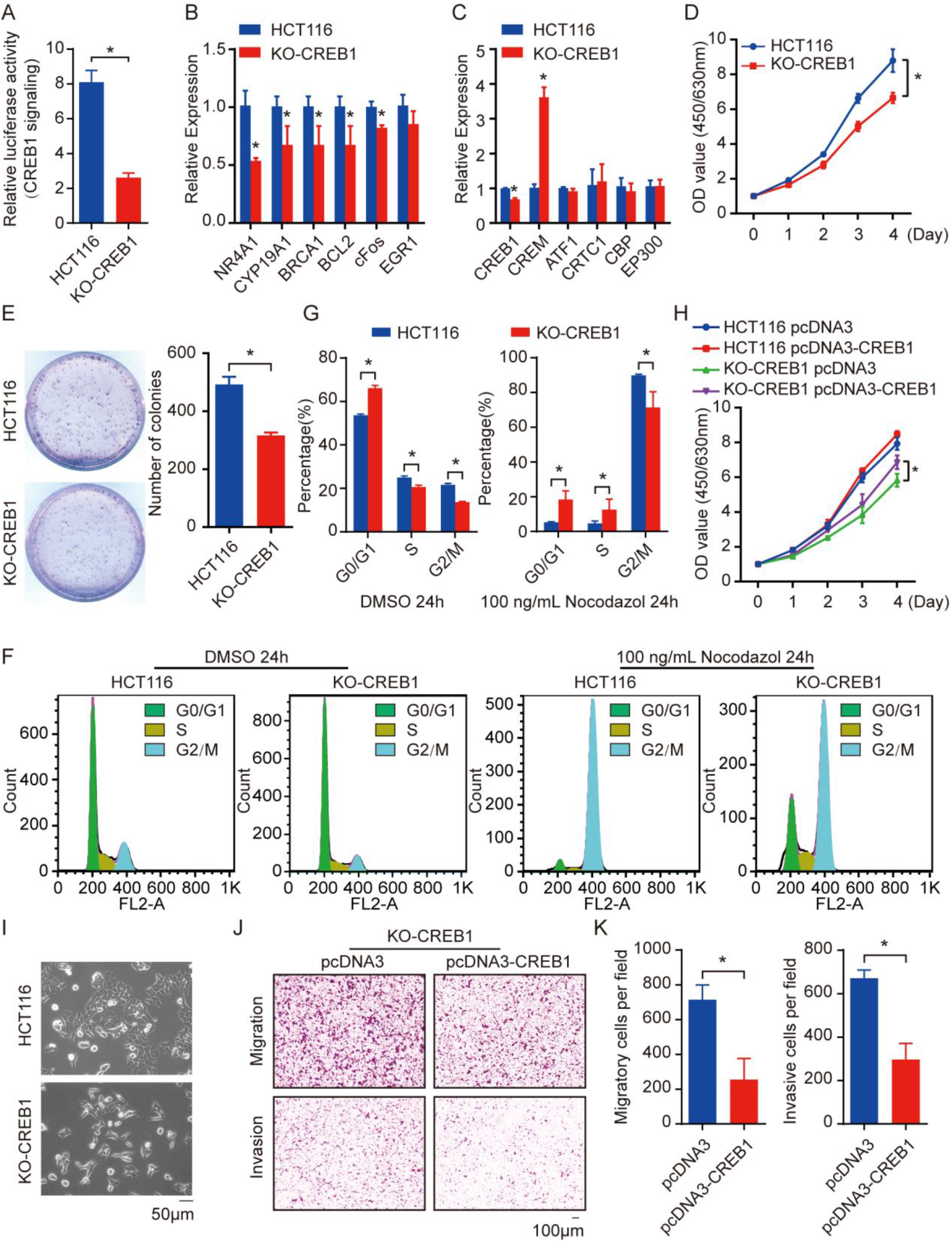
Effect of knocking out CREB1 on cell proliferation, cell cycle, migration and invasion of HCT116 cells *in vitro*. (A-C) CREB1 pathway reporter assay (A), CREB1 pathway target genes (B), and CREB1 pathway related genes (C) were assessed in CREB1 knockout HCT116 cells; wild-type HCT116 cells were used as vehicle control. (D-E) CCK-8 (D) and colony formation assays (E) showing the effects of CREB1 knockout on the proliferation of HCT116 cells *in vitro*. The quantification of colony formation assays is shown on the right panel. (F-G) Proportions of cells in each phase of the cell cycle were determined by flow cytometry analysis. Both HCT116 and KO-CREB1 cells were treated with DMSO or 100 ng/mL Nocodazole. Quantification of the percentage of cells in each phase of the cell cycle phase was determined for the plots and represented in a graph. (H) CCK-8 assay showing the effect of overexpression on the growth of HCT116 and KO-CREB1 cells. (I) Phase-contrast images of HCT116 and KO-CREB1 cells. (J-K) Restoration of CREB1 expression in KO-CREB1 cells is sufficient to inhibit the migratory and invasive ability of cells. Quantification of the cells was counted for plots and represented in a graph. All experiments were repeated at least three times with similar results. Error bars represent SEM. \**p* < 0.05, as assessed by Student’s *t*-tests.

Using CCK-8 and colony formation assays, we found that knockout of CREB1 significantly inhibited HCT116 cell proliferation (Figure 1D and 1E). Flow cytometry was used to assess whether the absence of CREB1 in HCT116 cells was associated with changes in the proportion of cells in each phase of the cell cycle. As shown in Figure 1F and 1G, the percentage of KO-CREB1 cells in G0/G1-phase was significantly increased compared with that of the wild-type HC116 cells during nocodazole treatment and untreated. In addition, we exogenously introduced CREB1 to confirm its function in cell proliferation and the cell cycle (Supplementary figure 1F). Consistent with the knockout experiments, rescue of CREB1 in KO-CREB1 cells resulted in enhanced cell proliferation and G1/S transition (Supplementary figure 1G). However, ectopic expression of CREB1 in wild-type HCT116 cells had no influence on cell proliferation or cell cycle progression (Figure 1H and Supplementary figure 1G). In addition, disruption of CREB1 in HCT116 cells did not lead to cell apoptosis or cell death (Supplementary figure 1H).

Upon detecting the phenotypic change of the KO-CREB1 cells, we found that the absence of CREB1 induced an obvious morphological change in the cells, from an epithelial-like shape to a spindle shape, meaning that CREB1 might contribute to cell motility (Figure 1I). We next studied whether CREB1 ablation was involved in promoting the metastatic phenotype in HCT116 cells. *In vitro* transwell assays showed a 2-fold increase in the migration and invasion of KO-CREB1 cells (Guo et al., 2016a). More importantly, overexpression of CREB1 in both wild-type and KO-CREB1 cells produced only a small number of migrating and invading cells (Figure 1J and 1K).

Taken together, these data demonstrate that disruption of CREB1 in HCT116 cells by the CRISPR/Cas9n system impairs cell proliferation by accumulating cells in the G0/G1 phase, but it increases migration and invasion, leading to remarkable EMT *in vitro*.

### Knockout of CREB1 alters multiple signaling pathways related to cell status

To further identify the gene pathways and molecular signatures that were altered upon knockout of CREB1, we conducted strand-specific RNA sequencing (ssRNA-seq) comparisons of wild-type and KO-CREB1 cells. Briefly, we identified 328 genes with significant differential expression (RSEM-EB-seq, 0.95 PPDE) between the wild-type and KO-CREB1 cells (Figure 2A, Supplementary table 1). In addition, GSEA confirmed that genes in the cAMP pathway were enriched in wild-type cells (Figure 2B).

**Figure 2.**
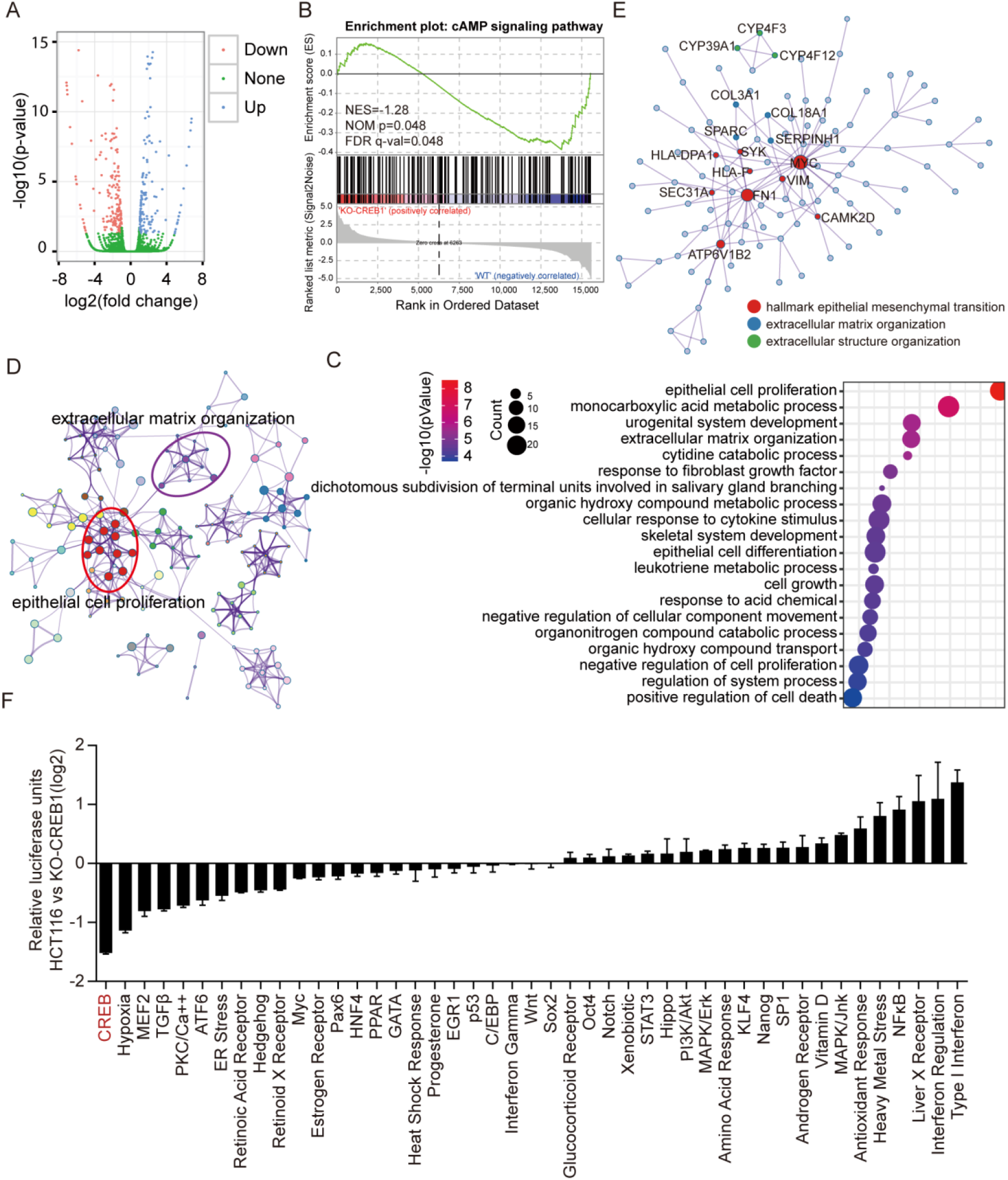
Knocking out CREB1 in HCT116 cells modulates multiple signaling pathways. (A) Volcano map indicating up-or downregulation of genes between wild-type and CREB1 knockout HCT116 cells. (B) GSEA revealing the enrichment of cAMP signaling pathway genes in CREB1 wild-type cells. Genes are ranked by changes in KO-CREB1 cells. (C-E) GO (C and D) and protein-protein interaction (E) analysis of the differentially expressed protein coding genes identified by comparing wild-type and CREB1 knockout HCT116 cells. (F) The 45-pathway assay analysis revealed the effects of CREB1 ablation on the activity of various pathways in HCT116 cells. The data were log2-transformed. All pathway reporter assays and qPCR experiments were performed at least three times. Error bars represent SEM. \**p*< 0.05, as assessed by Student’s *t*-tests.

To obtain a global overview of the biological functions of dysregulated genes upon knocking out CREB1, we performed functional annotation using Metascape analysis for Gene Ontology and Protein-protein Interaction (Tripathi et al., 2015). In addition to the GO terms and pathways previously associated with CREB1, such as “cell proliferation”, “positive regulation of transcription from RNA polymerase II promoter” and “TGF-beta signaling pathway”, we observed enrichment of genes involved in “extracellular matrix”, “cell adhesion”, “regulation of actin cytoskeleton” and “ECM-receptor interaction” (Figure 2C, 2D and 2E). Interestingly, these extracellular matrix-associated GO terms and pathways were in line with the role of CREB1 observed in the invasive KO-CREB1 cells.

Another method, 45-Pathway Reporter Arrays, was used to simultaneously assess 45 different signaling pathways in CREB1 knockout HCT116 cells. As expected, the CREB1 pathway exhibited the most prominent decrease when comparing knockout with wild-type HCT116 cells. More interesting, many other pathways involved in cell growth, such as hypoxia, MEF2, TGF-beta, and MYC, declined significantly in knockout cells. In contrast, INF, LXR, and NF-κB were largely upregulated, showing intrinsic immune and inflammatory activation (Figure 2F).

### CREB1 facilitates the G1/S transition by transactivation of CCAT1 and E2F1

To characterize the mechanism by which knockout of CREB1 induced G0/G1 arrest, we analysed the expression of 15 genes involved in the G1 phase and G1/S transition in wild-type HCT116 and KO-CREB1 cells. As shown in Figure 3A and 3B, E2F1, a gene known to be important in cell cycle progression, was decreased in KO-CREB1 cells, and the expression of cell cycle inhibitors, CDKN2B (p15), CDKN2A (p21) and CDKN1A (p27), was markedly induced in KO-CREB1 cells. Intriguingly, CCND1 (Cyclin D1) and CCNE1 (Cyclin E1), which have been reported to be transcriptionally regulated by CREB1 (Lee et al., 1999, D’Amico et al., 2000, Nagata et al., 2001, Boulon et al., 2002), did not change in either RNA or protein levels in KO-CREB1 cells.

**Figure 3.**
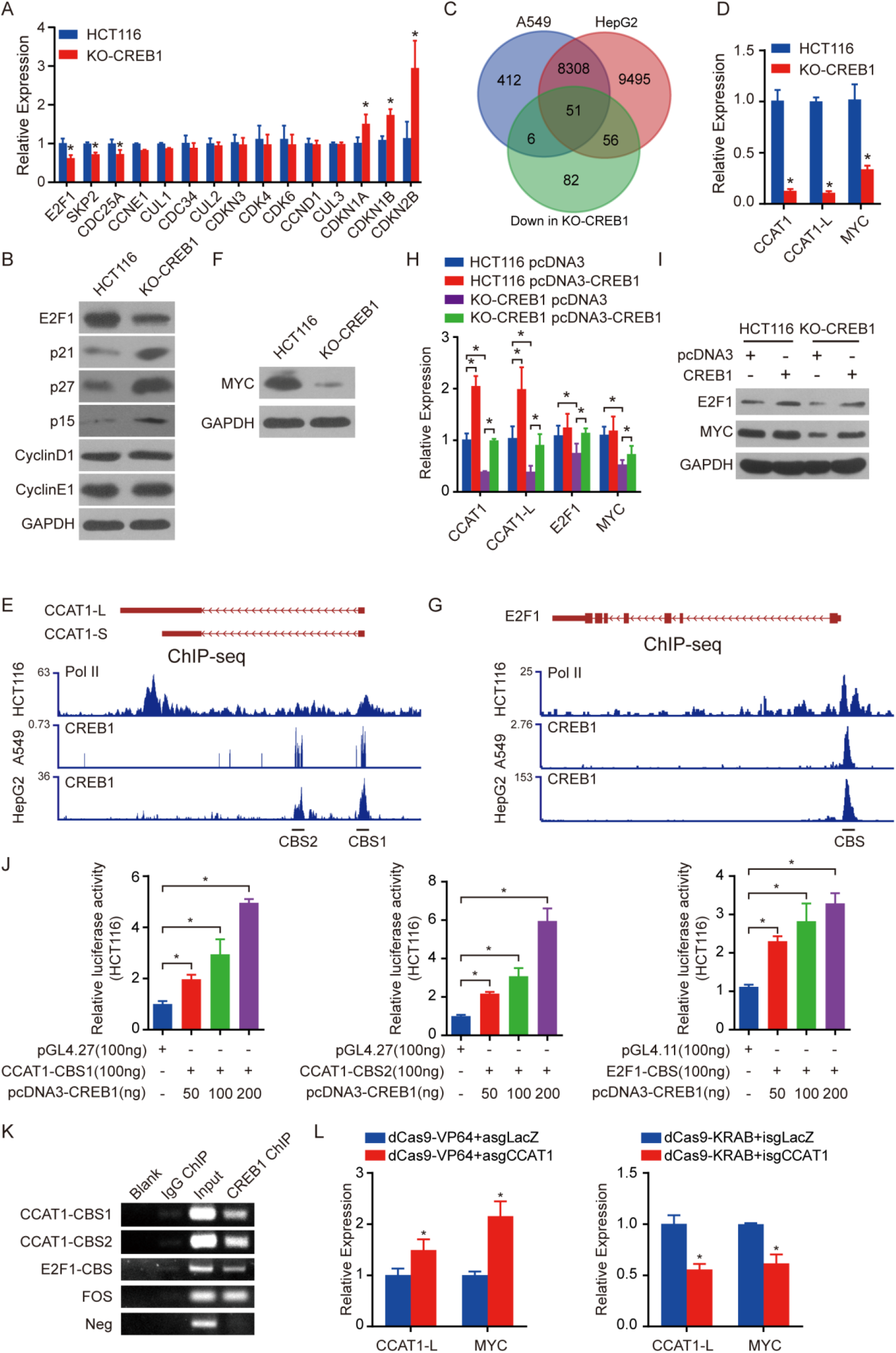
Downregulation of CCAT1 and E2F1 transcription by CREB1 ablation inhibits cell proliferation and the cell cycle. (A-B) qPCR (A) and Western blot (B) analyses of the expression of genes involved in the G1 phase and G1/S transition in HCT116 and KO-CREB1 cells. GAPDH was used as a loading control. (C) Venn diagram showing the proportion of CREB1-occupied genes whose expression is significantly downregulated in response to knockout of CREB1 in HCT116 cells. (D) qPCR analysis of the expression of CCAT1, CCAT1-L and MYC in HCT116 and KO-CREB1 cells. (E) Schematic representation of the CCAT1, and CCAT1-L (F) gene structures, RNA polymerase II levels of HCT116 cells, and CREB1 levels of HepG2 and K562 cells. (F) Western blot of MYC in HCT116 and KO-CREB1 cells. GAPDH was used as a loading control. (G) Schematic representation of the E2F1 gene structure, RNA polymerase II levels of HCT116 cells, and CREB1 levels in HepG2 and K562 cells. (H) qPCR analysis of CCAT1, CCAT1-L, E2F1 and MYC expression upon introduction of CREB1 in both HCT116 and KO-CREB1 cells. (I) Western blot analysis of E2F1 and MYC protein levels upon introduction of CREB1 in both HCT116 and KO-CREB1 cells. (J) Luciferase reporter assays showing the dose-dependent effect of CREB1 overexpression on CCAT1-CBS1, CCAT1-CBS2 and E2F1-CBS. (K) Semiquantitative ChIP-PCR assays demonstrating an *in vivo* interaction between CREB1 and the potential CREB1 binding site of CCAT1 and E2F1. The CREB1 binding site in the FOS gene promoter was used as a positive control, and the first intronic region of E2F1 was used as a negative control (Neg). (L) Quantitative gene expression analysis of CCAT1, CCAT1-L and MYC in HCT116 cells by dCas9-VP64 (CRISPRa) and dCas9-KRAB (CRISPRi) cotransfected with sgRNAs targeted upstream or downstream of the CCAT1 transcription start site, respectively. The data are representative of three independent experiments. Error bars represent SEM. \**p*< 0.05, as assessed by Student’s *t*-tests.

Because of CREB1’s central role in transcription, we compared genes whose expression was downregulated in KO-CREB1 cells with CREB1-occupied genes, using existing CREB1 ChIP-seq data from A549 and HepG2 cells (Figure 3C, Supplementary table 2). We observed that 51 genes were found in both the A549 and HepG2 datasets. From the 51 candidates, we noticed a lncRNA gene, CCAT1, that had previously been demonstrated to be important in several stages of CRC, especially in relation to cell proliferation (Figure 3D and 3E) (Kim et al., 2015). Moreover, MYC, which was positively regulated by CCAT1-L via its role in recruitment of CTCF in HCT116, was obviously downregulated in KO-CREB1 cells (Figure 3D and 3F) (Xiang et al., 2014). The promoter of E2F1 was also shown to be occupied by CREB1 in all four ChIP-seq datasets despite the absence of E2F1 in the downregulated genes from the RNA-seq experiments (Figure 3G). Additionally, CREB1 was found to have the potential to promote the transcription of MYC in light of CREB1 ChIP-seq (Supplementary figure 2A). Thus, these data suggest that CREB1 is involved in the transactivation of CCAT1 lncRNA and transcriptional regulatory proteins.

To test the above results and assumptions, rescue experiments were first performed, and the results showed that CREB1 overexpression in HCT116 cells and restoration in KO-CREB1 cells remarkably enhanced the expression of CCAT1, but it significantly increased MYC and E2F1 only in KO-CREB1 cells (Figure 3H and 3I). Then, we subcloned the CREB1 binding sites (CBSs) in CCAT1 (named CCAT1-CBS1 and CCAT1-CBS2) into pGL4.27, which contains a minimal promoter. E2F1-CBS1, which already contains the promoter of E2F1, was subcloned into the promoterless vector pGL4.11. The constructs were cotransfected into HCT116 cells with pcDNA3-CREB1. As expected, the CBS fragments responded to CREB1 in a dose-dependent manner (Figure 3J). Third, we performed a ChIP assay with HCT116 cells using an antibody against CREB1 and observed an enrichment of CCAT1 and E2F1 promoter fragments containing the CBS region over that of the normal IgG, which served as a nonspecific control (Figure 3K). In contrast, CREB1 did not occupy on the promoter of MYC in HCT116 cells (Supplementary figure 2B). Taken together, these results indicate that CCAT1 and E2F1, but not MYC, are transcribed by CREB1.

To further clarify the effect of dysregulation of CCAT1 on MYC, we performed *in cis* overexpression and downregulation of CCAT1-L in HCT116 cells based on CRISPRa and CRISPRi, respectively. Compared to the control, the overexpression of CCAT1-L indeed enhanced MYC expression (Figure 3L). Consistently, the downregulation of CCAT1-L reduced the expression of MYC (Figure 3L). Previous studies have shown that MYC acts as a transcriptional activator of E2F1 and CCAT1, but it acts as a repressor to CDKN1A, CDKN1B and CDKN2B. Then, we conducted MYC overexpression and knockdown assays to explore its role in these genes. As a result, MYC negatively regulated the expression of CDKN1A, CDKN1B and CDKN2B, but it did not alter E2F1 and CCAT1 (Supplementary figure 2C-F). These data suggest that CREB1 influences the expression of CCAT1, MYC and E2F1. Then, the downregulation of MYC was found to increase the expression of growth-arresting genes CDKN1A, CDKN1B and CDKN2B.

In summary, these results reveal that CREB1 depletion negatively impacts cell proliferation and G1/S transition by directly downregulating the transcription of the lncRNA CCAT1 and the transcription factor E2F1.

### Increasing the migration and invasion by CREB1-deficiency depends on NF-κB pathway activation

To explore the mechanism by which CREB1-deficiency contributes to increased cell migration and invasion, we measured the expression of EMT markers in wild-type and KO-CREB1 HCT116 cells. As shown in Figure 4A and 4B, knockout of CREB1 in HCT116 cells enhanced the expression of ZEB1, Vimentin, VIM-AS1 and Slug and concomitantly decreased E-cadherin levels (Boque-Sastre et al., 2015). Conversely, CREB1 overexpression in HCT116 cells and its restoration in KO-CREB1 cells increased the expression of E-cadherin and reduced Vimentin and Slug (Supplementary figure 4A).

**Figure 4.**
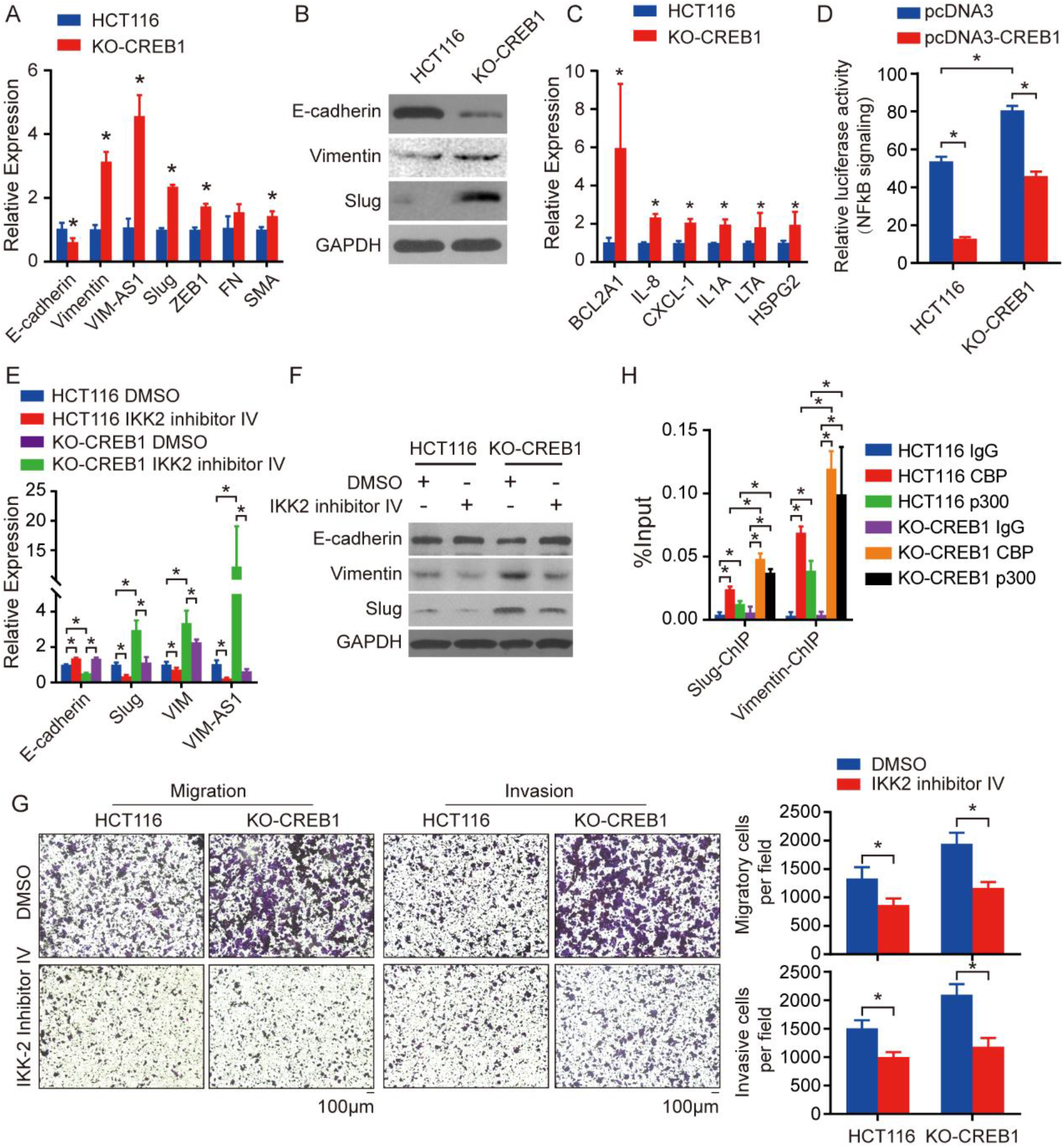
Knockout of CREB1 induces the EMT phenotype by activating the NF-κB pathway via competition of coactivators CBP/p300. (A-B) qPCR (A) and Western blot (B) analyses of EMT marker RNA and protein levels in HCT116 and KO-CREB1 cells. (C) qPCR analysis of NF-κB pathway target genes upon knockout of CREB1 in HCT116 cells. (D) NF-κB reporter assays upon CREB1 overexpression in HCT116 and restoration in KO-CREB1 cells. (E-F) qPCR (E) and Western blot (F) analyses of the expression of EMT markers after treatment of both HCT116 and KO-CREB1 cells with a NF-κB pathway inhibitor. (G) Transwell migration and invasion assays were performed to evaluate the migratory and invasive ability of both HCT116 and KO-CREB1 cells treated with NF-κB pathway inhibitors. Quantification of the cells was determined for plots and is represented on the right panel. (H) The regions of the p65-binding site of related EMT markers were quantified and were normalized to input levels following CBP and p300 ChIP-qPCR assays, which were performed on both HCT116 and KO-CREB1 cells. The data are representative of three independent experiments. Error bars represent SEM. \**p*< 0.05, as assessed by Student’s *t*-tests.

Our 45-pathway reporter assays revealed that the NF-κB signaling pathway, which is essential for EMT (Huber et al., 2004), was significantly upregulated in KO-CREB1 HCT116 cells. By qPCR analysis, the mRNA levels of the NF-κB signaling target genes BCL2A1, IL-8, and CXCL-1 were also significantly upregulated by the loss of CREB1 in HCT116 cells (Figure 4C) (Zong et al., 1999, Kunsch et al., 1994, Wood et al., 1995). The re-expression of CREB1 in KO-CREB1 cells reversed the induction of NF-κB signaling pathway (Figure 4D).

To further investigate whether the NF-κB signaling pathway is closely involved in CREB1 knockout-induced EMT, we used IKK-2 inhibitor IV, a selective inhibitor of the NF-κB signaling pathway, to repress its activity in HCT116 and KO-CREB1 cells. The NF-κB signaling reporter assay and the expression profile of its target genes confirmed the inhibition of the pathway in both HCT116 wild-type and KO-CREB1 cells (Supplementary figure 3B and 3C). Next, EMT markers were analysed by qPCR and Western blot in HCT116 and KO-CREB1 cells treated with 10 μM IKK-2 inhibitor IV for 6 h. As shown in Figure 4E and 4F, IKK-2 inhibitor IV treatment increased the expression of E-cadherin but decreased the levels of the mesenchymal marker. Consistently, inhibition of NF-κB pathway activity with IKK-2 inhibitor IV decreased the migration and invasion ability of HCT116 and KO-CREB1 cells (Figure 4G). Taken together, these results suggest that the NF-κB pathway promotes the EMT that is induced by CREB1 knockout in HCT116 cells.

Previous data suggest that the activity of the NF-κB pathway is positively correlated with the phosphorylation of p65; thus, the augmentation of the NF-κB pathway may be due to the increasing level of phosphorylation of p65 (Perkins and Gilmore, 2006). We then detected the protein levels of total and phosphorylated p65 as well as the expression of IKKa, IKKb and IkBa in wild-type and KO-CREB1 HCT116 cells. However, no obvious changes in the expression levels of these proteins were observed (Supplementary figure 3D). Thus, we hypothesized that other factors, such as CREB-binding protein (CBP) and its paralog p300, have a modulating role in the NF-κB signaling pathway independent of inducible nuclear translocation and posttranslational modifications (Gerritsen et al., 1997). To test this hypothesis, we detected the occupancy of CBP/p300 at the p65 binding site in the promoter of Slug and Vimentin with ChIP experiments for CBP and p300 in wild-type and KO-CREB1 HCT116 cells (Choi et al., 2015, Boque-Sastre et al., 2015, Chua et al., 2006). As shown in Figure 4H and Supplementary figure 3E, binding of CBP/p300 following CREB1 knockout was specifically enriched in the promoter of EMT markers (Slug and Vimentin) and the NF-κB signaling pathway target gene IL-8, indicating mutual competition of transcriptional coactivators between the two pathways.

Collectively, these experiments demonstrate that disruption of CREB1 releases CBP/p300, leading to the activation of the NF-κB signaling pathway and consequent transcription of EMT-associated genes, revealing a previously unknown mechanism for pathway activation mediated by CREB1 depletion.

## Discussion

High levels of CREB have been observed presented in many kinds of tumours, and it correlates with a high risk of tumorigenesis and a poor clinical outcome (Wu et al., 2007, Chhabra et al., 2007, Bulun et al., 2009, Shankar et al., 2005, Shankar and Sakamoto, 2004). In this report, we demonstrate that knockout of CREB1 impedes cell proliferation resulting in dysregulation of the G1/S phase transition, which is driven, at least in part, by the transcriptional downregulation of the lncRNA CCAT1 and of E2F1 (Figure 5, upper panel). However, We also observed increased cell migration and invasion in the setting of CREB1-deficiency, and this was probably due to the induction of EMT via NF-κB pathway activation (Figure 5, lower panel). Thus, our findings provide a mechanism underlying CREB1 contributes to tumour cell plasticity, which leads to a better understanding of the complexity of the CREB1 function and regulatory network.

**Figure 5.**
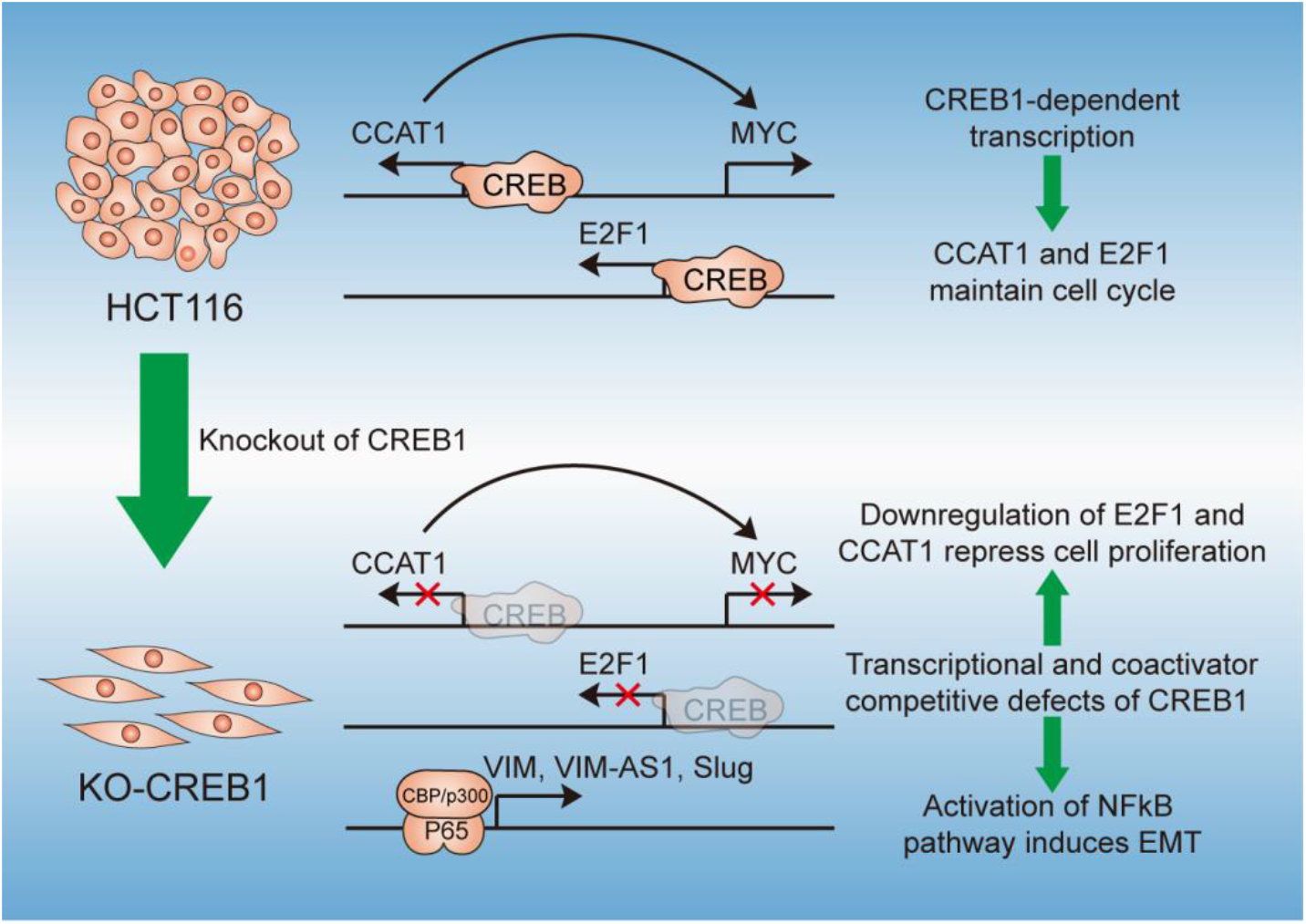
A functional model of CREB1 for controlling transition of tumour cells between proliferation and metastatic states by transactivation of lncRNA CCAT1 and competition of coactivators CBP/p300.

Given this complexity, we sought direct experimental demonstration that CREB1 can indeed promote cell proliferation as well as inhibit metastasis. Using CREB1 knockout cells, re-expression of CREB1 remarkably reduce the metastatic properties (Figure 1J and 1K). At the same time, exogenous CREB1 could increase proliferation in CREB1-deficency cells. Nevertheless, the proliferation of wild-type cells was unresponsive to the overexpression of CREB1, we reported previously that the metastatic ability of the cells was greatly reduced upon forced expression of CREB1 (Guo et al., 2016b). In addition, the regulation of MYC by CREB1/CCAT1 axis did not contribute to the expression of ITGB and ITGAV, which are involved in the motility and invasiveness of breast cancer cells (Supplementary figure 3F) (Hong et al., 2012). Thus, the opposite role of oncogenes that can promote proliferation pathways but simultaneously suppress migration pathways might be an intrinsic mechanism of cancer cell plasticity in tumours, although more examples need to be uncovered.

In this study, the lncRNA CCAT1 and the cell cycle-associated transcription factor E2F1 were validated as a novel ncRNA target and protein target of CREB1, respectively. It is worth noting that CCAT1, which was identified as an upregulated lncRNA in CRC, was associated with the malignant progression of CRC, involving cell proliferation, invasion and migration *in vitro* and *in vivo* (Nissan et al., 2012, Kam et al., 2013, Xin et al., 2016). Although overexpression of CREB1 in HCT116 cells can promote the expression of CCAT1, as well as CCAT1-L, by approximately 2-fold, the restoration of CREB1 in HCT116 KO-CREB1 cells can increase CCAT1 levels, approaching but not exceeding its expression in HCT116 cells with wild-type CREB1 (Figure 3H). In addition, the expression of E2F1 was not changed significantly in HCT116 cells with forced expression of CREB1. Thus, CREB1 may only contribute to basal transcription of CCAT 1 and E2F1, and it may maintain cell proliferation through basal control of the cell cycle, especially in the G1/S phase transition. Additionally, CREB1 has been reported to regulate the cell cycle by transactivation of its downstream genes, including CCND1, CCNE1, CCNA1 and CCNA2, while the expression of CCND1 and CCNE1 did not change significantly in the RNA or protein level in CREB1-knockout HCT116 cells. The CREB1 occupancy of the CRE motif in the CCND1 and CCNE1 genes was validated with ChIP-PCR, suggesting a direct role for CREB1 in regulation CCND1 and CCNE1 in HCT116 cells (Supplementary figure 3H). By studying genomic sequences, the complex promoter of the CCND1 and CCNE1 genes were found to contain many transcription factor binding sites that may control their transcriptions depending on the particular cellular or functional context (data not shown). Therefore, the absence of CREB1 in HCT116 cells might be compensated for by other genes.

Our finding that CREB1 overexpression leads to inhibition of cell migration and invasion through suppression of the NF-κB pathway supports notion that the NF-κB pathway function in angiogenesis, invasion and metastasis (Chua et al., 2006, Huang et al., 2001, Huber et al., 2004). However, none of previous reports has linked the effect of CREB1 on NF-κB pathway activity to cell motility and invasiveness, and none has implicated the EMT markers in the cellular changes elicited by CREB1. Our results demonstrate that the disruption of CREB1 leads to a loss of its ability to compete with p65 for limiting amounts of coactivators CBP/p300, which in turn increases properties that are essential for metastasis. These data, however, do not mean that CREB1 affects invasion and metastasis only through regulation of the NF-κB pathway-mediated EMT. The fact that CREB1 has been reported to induce the expression of MMP-2/9 and proteins associated with EMT in Renal cell carcinoma (Wang et al., 2017). Nevertheless, as inhibition of the NF-κB pathway activity by IKK-2 inhibitor IV is sufficient to counteract the repressive effect of CREB1 in migration and invasion, it is apparent that a primary effect of CREB1 on motility in HCT116 cells is mediated through its inhibition of the NF-κB pathway.

Ultimately, the findings reported here raise the possibility that inhibition of CREB1 in human tumours might at times be contraindicated because its suppression may indeed promote metastasis and therefore, inhibition of the NF-κB pathways should be taken into consideration as a combinatorial treatment in therapeutic strategies involving CREB1.

## Materials and methods

### Key resources table

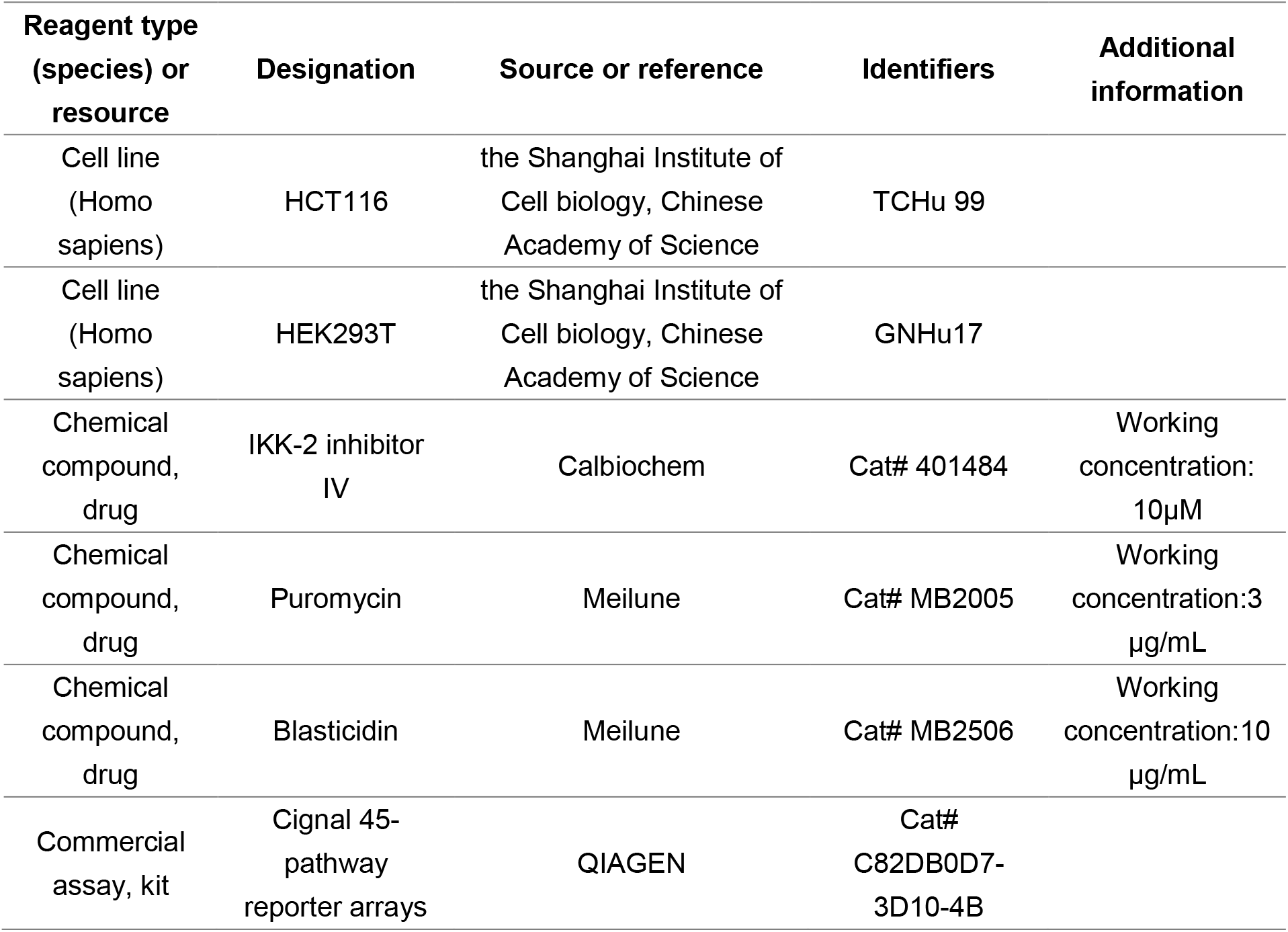
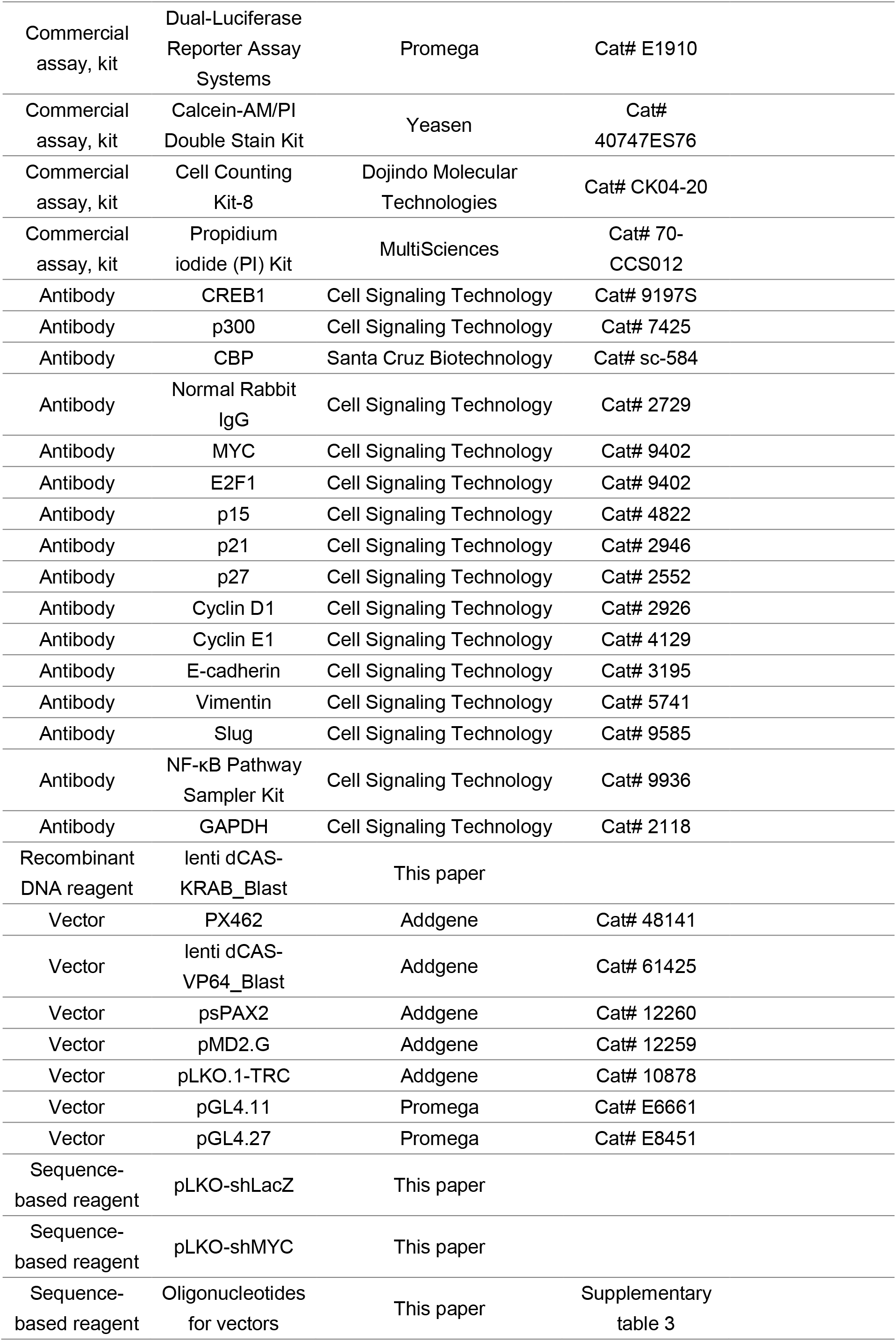
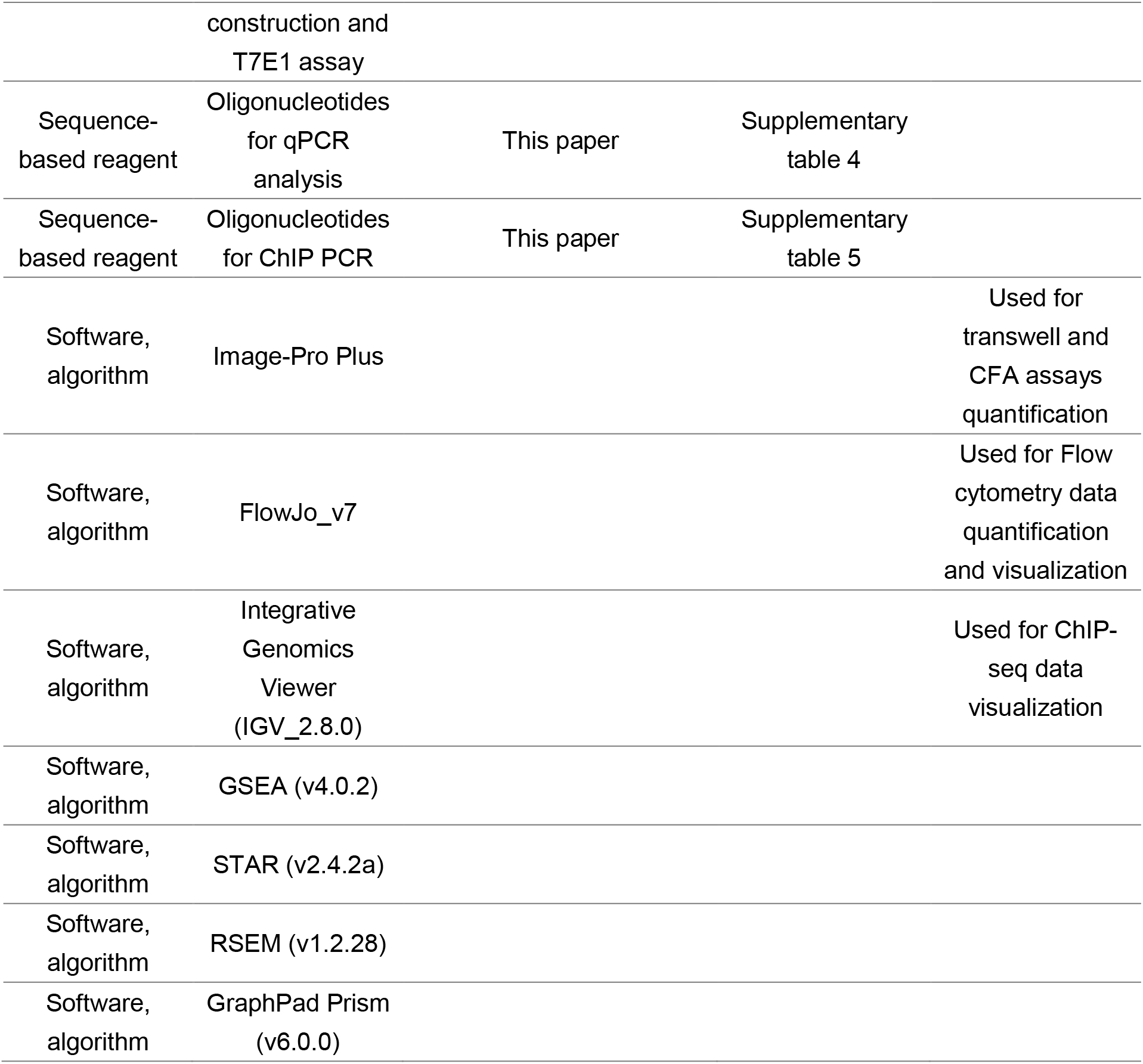

### Cell cultures and treatments

The human colon cancer cell line HCT116 and human embryonic kidney cell line HEK293T were obtained from the Shanghai Institute of Cell biology, Chinese Academy of Science. Cell lines were maintained using standard media and conditions. To inhibit the NF-κB pathway, HCT116 and KO-CREB1 cells were treated with different concentrations of 5, 10, 15 and 20 μM IKK-2 inhibitor IV (Calbiochem, #401484) for 6 hr in a serum-free medium, and DMSO (Sigma) was used as a control.

### Cell transfection

Transfection of plasmid DNA was performed using Lipofectamine LTX reagent or Lipofectamine 2000 (Invitrogen), and all RNA transfections were performed at a final concentration of 50 nM using Lipofectamine 2000 (Invitrogen) according to the manufacturer’s instruction. Besides, Cignal 45-pathway reporter arrays plasmids were transfected with ViaFect™ Transfection Reagent (Promega).

### Knockout of CREB1

The CRISPR/Cas9 Nickase (CRISPR/nCas9) system was used to construct the HCT116 KO-CREB1 cells (Guo et al., 2016a). The HCT116 cells were transfected with PX462-C1C2 using Lipofectamine 2000 and cultured in the presence of puromycin (3 μg/mL, Sigma) for 48 hr after post-transfection 24 hr. Single-cells clones were generated by dilution assays and tested for knockout of CREB1 expression by Western blot.

### RNA-seq sequencing and analysis

RNA-sequencing was performed by RiboBio Co., Ltd. Briefly, ribosomal RNA was depleted from HCT116 wild-type and KO-CREB1 cells by using a Ribo-Zero rRNA Removal Kit (Epicentre). Then, the RNA was used as input for the Illumina TruSeq® Stranded Total RNA HT/LT Sample Prep Kit (Illumina). Libraries were pooled and sequenced on an Illumina Hiseq 3000 PE150 platform. A minimum of 50 M reads were generated per sample. RNA-sequencing reads were trimmed for adaptor sequence and mapped to the GENCODE V19 using STAR and RSEM. Then, protein coding genes exhibiting differential expression with PPDE>0.95 were included in Gene Ontology and Protein-protein Interaction analyses using Metascape (Tripathi et al., 2015). For GSEA, the cAMP signaling pathway gene set was generated according to the gene list from KEGG; then normalized expression data were analyzed and visualized with GSEA software (version 3.0, http://www.broadinstitute.org/gsea/) (Subramanian et al., 2005, Kanehisa et al., 2018).

### ChIP-seq Data analysis and visualization

Both RNA polymerase II and CREB1 ChIP-seq data were previously published and were obtained from NCBI GEO accession numbers: GSM935426(HCT116, RNA polymerase II), GSM1010808(HepG2, CREB1) and GSM1010719(A549, CREB1). Peaks were identified using MACS2 with default parameters (Feng et al., 2012). All data were visualized using Integrative Genomics Viewer (IGV) software.

### Vectors construction

The pcDNA3 vector (Invitrogen) was used for generation CREB1 and MYC expression vectors. CBS1 and CBS2 of CCAT1 and E2F1-CBS upstream of E2F1 were amplified from HCT116 genome and inserted into pGL4.27 and pGL4.11 (Promega) in the sense orientation relative to the luciferase coding sequence, respectively. For stable expression of dCas9-KRAB, the lenti dCAS-KRAB_Blast vector was modified from lenti dCAS-VP64_Blast vector (Addgene) as follows: the KRAB domain (from ZNF10) was amplified by PCR from HEK293T cDNA and cloned into lenti dCAS-VP64_Blast vector using *Bam*HI and *Bsp*1407I double digestion by ClonExpress MultiS kit (Vazyme). For transient expression of sgRNA, the pU6-sgRNA vectors was constructed by subcloning the U6 promoter-sgRNA scaffold from PX462 (Addgene) to pMD-18T vector (Takara). The sgRNA expression plasmids were cloned by insert annealed oligos into the pU6-sgRNA vector that was digested by *Bbs* I (NEB). shRNA for *LacZ* (negative control) and MYC were generate according to the pLKO.1 protocol from Addgene. Cignal 45-pathway reporter arrays, the CREB pathway reporter and the NF-κB pathway reporter that measure the activity of the corresponding pathway were purchased from Qiagen. The primers used for vector construction are listed in Supplementary table 3.

### Genomic DNA extraction, amplification and T7E1 assay

Genomic DNA (gDNA) from HCT116 single-cells clones, which transfected with sgRNAs targeted CREB1 were extracted with Hipure Tissue DNA mini kit (Magen) according to the manufacturer’s instruction. Then, the target region was amplified by Phusion® High-Fidelity DNA Polymerase (NEB), and the indel of the sgRNAs was assessed by the T7 endonuclease I assay (Viewsolid Biotech). The primers used for T7E1 assay are listed in Supplementary table 3.

### Lentiviral Transduction for Stable cell lines

The lenti dCAS-VP64_Blast, lenti dCAS-KRAB_Blast, pLKO-shLacZ or pLKO-shMYC vectors were co-transcripted with packaging vectors psPAX2 and pMD2.G into HEK293T cells for lentivirus production. To establish stable cell lines, HCT116 cells were transduced by using the above lentiviral particles with polybrene (6 μg/mL, Sigma). After 24 hr, the infected cells were subjected to selection in medium containing blasticidin (10 μg/ml, Meilune) or puromycin (3 μg/ml, Meilune) for 5 days.

### RNA extraction and qPCR assays

Total RNA was extracted from HCT116 wild-type and KO-CREB1 cells with TRIzol reagent (Invitrogen) according to the manufacturer’s instruction. and then reverse-transcribed to cDNA using the primescript™ RT reagent kit (Takara). Real-time PCR was carried out using SYBR Premix ExTaq™ (Takara) according to manufacturer’s instructions. GAPDH were employed as endogenous controls. The comparative Ct Method (ΔΔCT Method) was used to determine the expression levels of genes. All primer sets were purchased from Synbio Technologies (Suzhou, China), and the primer sequence used for qRT-PCR are show in Supplementary table 4.

### Cell viability assay

Cell viability was measured using Cell Counting Kit-8 (CCK-8, Dojindo Molecular Technologies). HCT116, KO-CREB1 and 24 hr post-transfection cells were seeded into a 96-well plate at 2×10^3^ cells per well. Then evaluating cell viability with CCK-8 at the time of 24, 48, 72, 96, 120 hr. The detected wavelength 450 nm, and the reference wavelength 630 nm were measured after incubation for 2 hr at 37°C.

### Calcein-AM/PI double staining

Calcein-AM/PI double staining of wild-type and KO-CREB1 HCT116 cells were performed according to the manufacturer’s instruction (Calcein-AM/PI Double Stain Kit, Yeasen). Brife, cells incubated in 1×Assay Buffer containing 0.67 μM Calcein-AM and 1.5 μM propidiumiodide (PI) at 37 °C for 15 min. After washing with PBS, cells were viewed in a fluorescence microscopy. The cells were counted using a Image Pro-Plus software.

### Cell cycle analysis

Cell cycle distribution was assessed by flow cytometry using Propidium iodide (PI) Kit (MultiSciences). HCT116, KO-CREB1 and 24 hr post-transfection cells were treated with vehicle or Nocodazole for 24 hr. Then the cells were resuspended in trypsin and centrifuged at 400 × g for 3 min at room temperature. The cell pellet was washed once by PBS and resuspended in 1 ml reagent A and in 10 μl reagent B. After incubating for 30 min at room temperature, the cell cycle was examined by flow cytometry.

### Colony formation assay

For evaluating effect of CREB1 knockdown on clonogenicity, cells were plated into 6-well plates with 2 × 10^3^ cells per well. After one week, colonies (>50 cells per colony) were fixed and stained with crystal violet in methanol. The colonies were counted using a Image Pro-Plus software.

### Luciferase reporter assays

Both forward and reverse transfection methods have been applied for luciferase reporter assay. And 45-pathway assay is performed with reverse transfection, forward transfection is employed for promoter assay, and pathway assay. For forward transfection, cells are plated in 48-well plate 24 hr before transfection. 48 hr post-transfection, cell extracts were assayed for luciferase activity using the Dual-Luciferase Reporter Assay Systems (Promega) according to the manufacturer’s instruction. Reverse transfection involves plating cells into the transfection complexes in a one day procedure. For promoter assay and 45-pathway assay, the Renilla luciferase was used as a control to normalize luminescence levels.

### Migration and invasion assays

For HCT116 and KO-CREB1 migration and invasion assay, the cells were suspended in 200 μl medium without FBS, and then were seeded into upper chamber of transwell inserts (8 μM pore size, Costar) with or without coated Matrigel (R&D System). The lower chamber of the transwell was filled with 750 μl medium supplemented with 10% FBS which function as a chemoattractant. After 24 hr incubation in 37°C, cells that migrated or invaded to the lower surface of the insert membrane were fixed in methanol and stained by 0.1% crystal violet. For IKK-2 Inhibitor IV treated cells migration and invasion assay, the cells were treated with 10 μM IKK-2 Inhibitor IV for 6 hr after 48 hr post-seeding. Then the cells were placed into the upper chamber with the non-coated or Matrigel-coated membrane, which was diluted with 200 μl serum-free culture medium. After the cells were incubated for 24 hr at 37°C in a humidified incubator, the cells adhering to the lower surface were fixed in methanol and stained by 0.1% crystal violet. The numbers of cells that have migrated or invaded were captured under Zeiss AxioObserver.Z1 (Carl Zeiss) at a magnification of 50×. And cell counting was processed by Image Pro-Plus software.

### Western blot analysis

Total protein was extracted using either TRIzol reagent (Invitrogen) as recommended or RIPA buffer (50mM Tris-HCl,pH 7.4, 150mM NaCl, 0.1% SDS, 1% Sodium deoxycholate, 1% Triton X-100, 2mM EDTA and 1×Protease Inhibitor Cocktail). Protein precipitation isolated by TRIzol was lysed in the buffer containing 1% DTT, 4% CHAPS, 7 M urea, 2 M thiourea and 2% ampholine. Equal total protein extracts were loaded and separated by SDS-polyacrylamide gel electrophoresis (SDS-PAGE), transferred onto Protran™ nitrocellulose membranes (Whatman). The following antibodies were used for western blot: CREB1 (CST, 9197S), MYC(CST, 9402S), E2F1(CST, 9402S), p15(CST,4822S), p21(CST, 2946S), p27(CST, 2552S), Cyclin D1(CST, 2926S), Cyclin E1(CST, 4129S), E-cadherin (CST, 3195S), Vimentin (CST, 5741S), Slug(CST, 9585), NF-κB Pathway Sampler Kit (CST, 9936) or GAPDH (CST, 2118S). GAPDH was used as the loading control.

### ChIP assay

HCT116 cells were cross-linked with 1% formaldehyde (Sigma) in new culture medium for 10 min and the reaction quenched with addition of glycine to a final concentration of 0.125 M at room temperature for 5 min. Cells were rinsed with cold PBS twice and scraped from the dishes. Pellet cells were resuspended in cell lysis buffer (20mM Tris-HCl at pH 8.0, 85 mM KCl, and 0.5% NP-40) and incubated on ice for 15 min, vortexing the cell suspension briefly every 5 min to facilitate the release of the nuclei. Nuclear lysis buffer (10mM Tris-HCl at pH 7.5, 1% NP40, 0.5% deoxycholate and 0.1% SDS) containing protease inhibitor mixture (Complete Mini tablets, Roche Diagnostics) was used to lyse the nuclei at a concentration of 1×107 cells/ml. After sonication, the chromatin lysates were divided into 500 μl aliquots. Each 500 μl sonicated chromatin (1×106 cells) pre-cleared with 10 μl Protein G Magnetic bead (Invitrogen). 5 μl (1%) of the supernatant was reserved as Input. Cleared samples were immunoprecipitated overnight in IP buffer with 10 μL of CREB1 (CST, 9197S), 10 μL of CBP (CST, 7425S), 10uL of p300 (Santa Cruz, sc-584) or 2μL negative control antibody rabbit IgG (CST, 2729). Immune complexes were successively washed twice with Low Salt Wash Buffer (0.1% SDS, 1% Trition X 100, 2mM EDTA, 20mM Tris-HCl (pH 8.0), 150mM NaCl), twice with High Salt Wash Buffer (0.1% SDS, 1% Trition X 100, 2mM EDTA, 20mM Tris-HCl (pH 8.0), 500mM NaCl), twice with LiCl Wash Buffer (0.25M LiCl, 1% NP-40, 1% deoxycholate, 1mM EDTA and 10mM Tris-HCl (pH 8.0)) and twice with TE Buffer (10mM Tris-HCl (pH 8.0) and 1mM EDTA) for 3 min. The precipitated chromatin was eluted and reversed cross-linked in ChIP Elution Buffer (1% SDS and 0.1M NaHCO3) containing proteinase K and RNase A two hours at 65 °C. The DNA was recovered and purified using Qiagen PCR purification kit. The semi-quantitative PCR or qPCR was performed to analyse the immunoprecipitated DNA. Primers are listed in Supplementary table 5.

### Statistical Analysis

Quantitative data were presented as the mean ± the standard error of the mean (SEM) from a minimum of three independent experiments. Comparisons between two groups were analyzed using the Student’s *t*-tests with n=3, unless otherwise indicated. Statistical analyses were performed with GraphPad Prism 6 (GraphPad Software Inc.). *p*<0.05 was considered to be statistically significant.

## Additional information

### Author contributions

L.H.Q., J.H.Y., H.Z. and B.L. conceived and designed the entire project. L.H.Q., J.H.Y. and H.Z. supervised the research. B.L., L.S.Z., C.M.Z., Q.J.H., Y.H.G., L.Q.W., P.Y., S.R.L., Q.L., Y.X.L. and J.H.Y. performed the experiments and/or data analyses. B.L. performed the genome-wide or transcriptome-wide data analyses. B.L., L.H.Q. and J.H.Y. contributed reagents/analytic tools and/or grant support. B.L., L.S.Z., C.M.Z., Q.J.H., Y.H.G., L.Q.W., P.Y., S.R.L., Q.L., Y.X.L., H.Z., J.H.Y. and L.H.Q. wrote and revised the paper. All authors discussed the results and commented on the manuscript.

## Acknowledgments

pSpCas9n(BB)-2A-Puro (PX462) was a gift from Feng Zhang (Addgene plasmid #48141). lenti dCAS-VP64_Blast was a gift from Feng Zhang (Addgene plasmid #61425). pLKO.1-TRC cloning vector was a gift from David Root (Addgene plasmid #10878). psPAX2 was a gift from Didier Trono (Addgene plasmid #12260). pMD2.G was a gift from Didier Trono (Addgene plasmid #12259).

## Additional files

**Supplementary figure 1.** Knockout of CREB1 in HCT116 cells.

**Supplementary figure 2.** CREB1 facilitates the G1/S transition by transactivating of CCAT1 and E2F1.

**Supplementary figure 3.** Increases in the migration and invasion of KO-CREB1 cells are due to the induction of EMT via activation of the NF-κB pathway.

**Supplementary table 1**. Genes differentially expressed upon knocking out of CREB1 in HCT116 cells identified by the RNA-seq analysis.

**Supplementary table 2**. The gene lists used for comparing ChIP-seq and RNA-seq.

**Supplementary table 3.** Primers used for vectors construction andT7E1 assay.

**Supplementary table 4.** Primers used for reverse transcription and real-time PCR

**Supplementary table 5. Primers used for ChIP PCR**

