## Supplementary figures for "CREB1 contributes colorectal cancer cell plasticity by regulating lncRNA CCAT1 and NF-κB pathways"

### Supplemental Figures

#### Supplementary figure 1

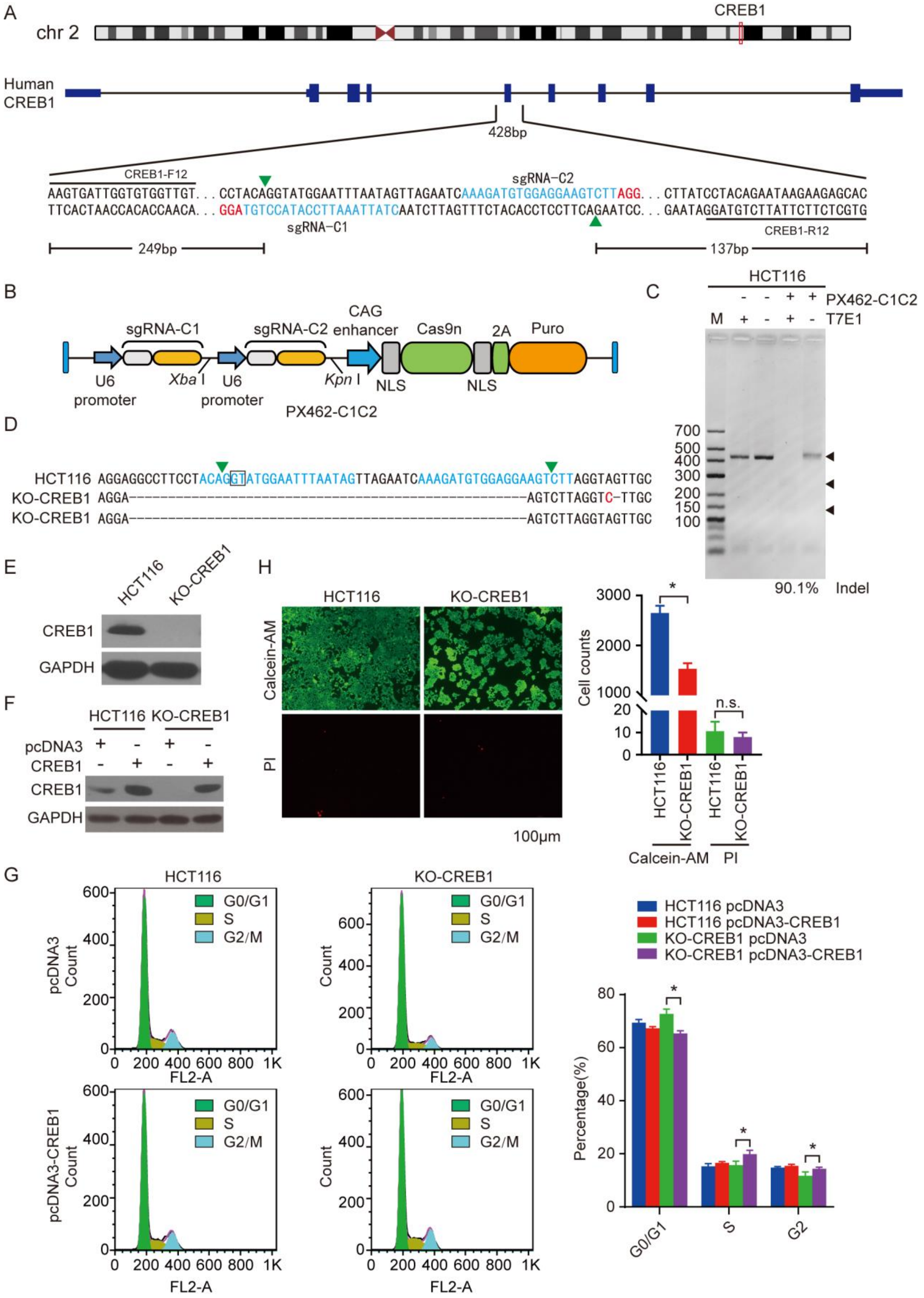


**Supplementary figure 1. Knockout of CREB1 in HCT116 cells.** (A) Cas9n target sites in exon 5 of the human CREB1 gene, positions of PCR primers used for amplification, and sizes of T7E1 cleavage products. Cas9n cleavage sites are highlighted by green triangles and the PAM motif is indicated by red font. The sgRNA target sites are indicated by blue. (B) A schematic of the tricistronic Cas9n CREB1 targeting vector. (C) Agarose gel results show T7E1 cleavage assays on PCR products amplified from the CREB1 locus. The substrate (428 bp) and two T7E1 cleavage products (249 bp and 137 bp) are indicated on the left by black triangles. The indel is indicated below the lane. (D) Representative indel sequences of the human CREB1 locus targeted by Cas9n. (E) Western blot analysis of CREB1 in HCT116 and KO-CREB1 cells. The experiment was repeated three times, and a representative blot is displayed. GAPDH was used as a loading control. (F) Western blot analysis of CREB1 in wild-type HCT116 and KO-CREB1 cells, where CREB1 was overexpressed and reintroduced, respectively; GAPDH served as a loading control. (G) The effect of CREB1 on the cell cycle in HCT116 and KO-CREB1 cells. Quantification of the percentage of cells in each phase of the cell cycle was determined for the plots and is represented in the right panel. Experiments were performed in triplicate. (H) Representative fluorescence micrographs of HCT116 and KO-CREB1 cells stained with Calcein-AM (green) and PI (red). Quantification of the cells was counted for plots and represented on the right panel. Error bars represent SEM. *$p < 0.05$, as assessed by Student's *t*-tests.

**Supplementary figure 2**

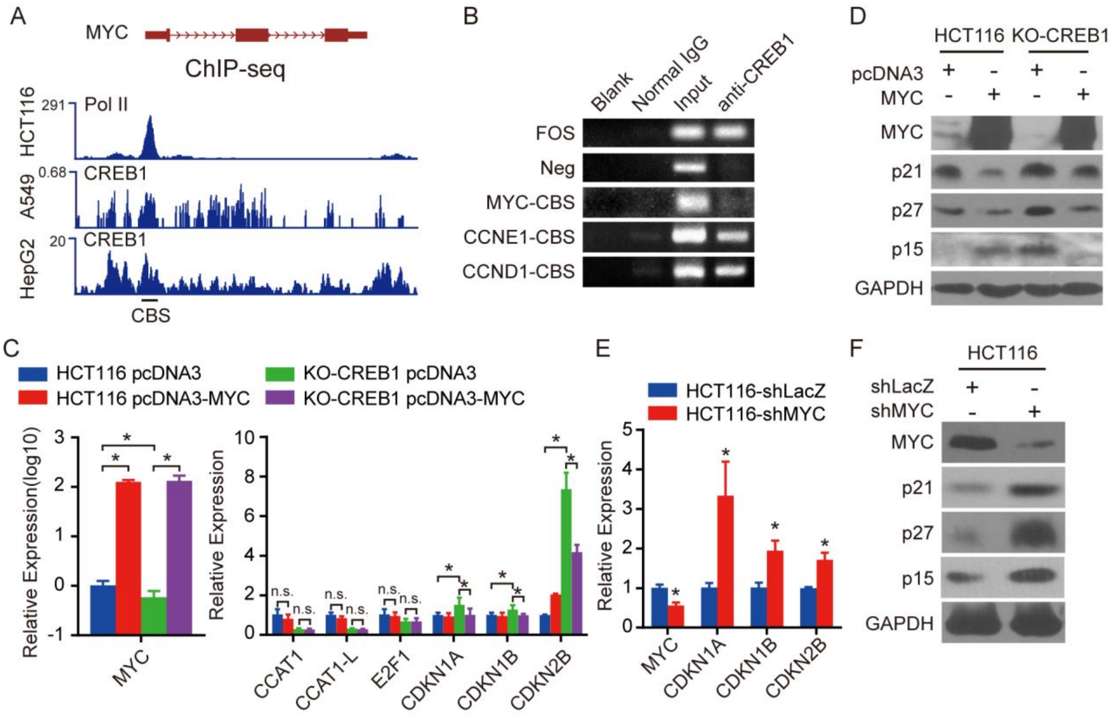


**Supplementary figure 2. CREB1 facilitates the G1/S transition by transactivating of CCAT1 and E2F1.** (A) Schematic representation of the MYC gene structure, RNA polymerase II levels of HCT116, and CREB1 levels of A549 and HepG2 cells. (B) Semiquantitative ChIP-PCR assays demonstrating an *in vivo* interaction between CREB1 and the potential CREB1 binding site of MYC, CCND1 and CCNE1. The CREB1 binding site in the FOS gene promoter was used as a positive control, and the first intronic region of E2F1 was used as a negative control (Neg). (C) qPCR analysis of MYC, CCAT1, CCAT1-L, E2F1, CDKN1A, CDKN1B and CDKN2B expression in MYC-overexpressing HCT116 and KO-CREB1 cells. (D) The levels of MYC, p15, p21 and p27 were determined by immunoblotting in MYC-overexpressing HCT116 and KO-CREB1 cells. (E-F) qPCR (E) and Western blot (F) analyses of MYC, CDKN1A, CDKN1B and CDKN2B expression in HCT116 cells that stably expressed either shLacZ (negative control) or shMYC .The results are representative data from three independent experiments. Error bars represent SEM. *$p< 0.05$, as assessed by Student's *t*-tests.

**Supplementary figure 3**

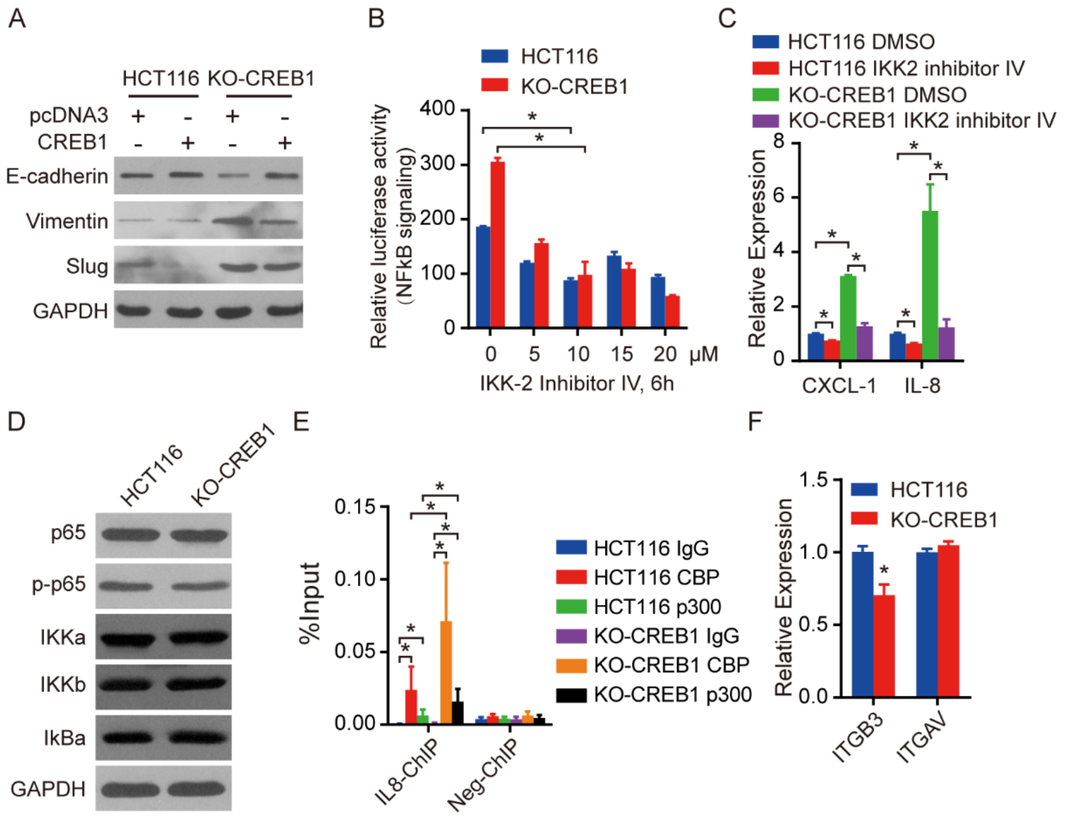


**Supplementary figure 3. Increases in the migration and invasion of KO-CREB1 cells are due to the induction of EMT via activation of the NF-κB pathway.** (A) Western blot analysis of EMT markers in both HCT116 and KO-CREB1 cells upon introduction of CREB1. (B) The NF-κB signaling reporter assay in HCT116 and KO-CREB1 cells treated with/without IKK-2 inhibitor IV at different concentrations. (C) qPCR analysis of NF-κB signaling pathway target genes in HCT116 and KO-CREB1 cells treated with 10 μM IKK-2 inhibitor IV for 6 h. (D) Western blot analysis of the protein levels of total and phosphorylated p65 as well as the expression of IKKa, IKKb and IkBa in HCT116 and KO-CREB1 cells. (E) The regions of the p65-binding site in the NF-κB signaling pathway target gene IL8 or nonspecific sites were quantified and normalized to input levels following CBP and p300 ChIP-qPCR assay in both HCT116 and KO-CREB1 cells. (F) qPCR analysis of ITGB3 and ITGAV in HCT116 and KO-CREB1 cells. The data are representative of three independent experiments. Error bars represent SEM. *$p < 0.05$, as assessed by Student's *t*-tests.
